# The neonatal microenvironment programs conventional and intestinal Tbet^+^ γδT17 cells through the transcription factor STAT5

**DOI:** 10.1101/658542

**Authors:** Darshana Kadekar, Rasmus Agerholm, John Rizk, Heidi Neubauer, Tobias Suske, Barbara Maurer, Monica Torrellas Viñals, Elena Comelli, Amel Taibi, Richard Moriggl, Vasileios Bekiaris

## Abstract

Interleukin(IL)-17-producing RORγt^+^ γδ T (γδT17) cells develop in the embryonic thymus and participate in type 3 immune responses. Herein we show that γδT17 cells rapidly proliferate within neonatal lymph nodes and gut, where upon entry they uniquely upregulate Tbet and co-express IL-17, IL-22 and interferon(IFN) γ in a STAT3 and retinoic acid dependent manner. Neonatal expansion was halted in mice conditionally deficient in STAT5 and its loss resulted in γδT17 cell depletion from all adult organs. Hyperactive STAT5 mutant mice showed that the STAT5A homologue had a dominant role over STAT5B in promoting γδT17 cell expansion and downregulating gut-associated Tbet. In contrast, STAT5B preferentially expanded IFNγ-producing γδ populations. Importantly, mice lacking γδT17 cells due to STAT5 deficiency displayed a profound resistance to experimental autoimmune encephalomyelitis. Our data identify for the first time STAT5 as a key molecular checkpoint allowing γδT17 cells to pass through a critical neonatal developmental window to acquire tissue-specific characteristics essential for infection and autoimmunity.

## Introduction

Interleukin(IL)-17 producing gamma delta (γδ) T cells (γδT17) are one of the major type 3 innate lymphocytes in the mouse, occupying the skin and most mucosal surfaces as well as secondary lymphoid organs. Their ability to constitutively produce IL-17 and to respond rapidly to cytokines like IL-7, IL-23 and IL-1β renders them as a critical part of innate immunity to infections (Cho et al., 2010; Conti et al., 2014) but also makes them highly pathogenic in a number of inflammatory models (Bekiaris et al., 2013; Michel et al., 2012; Sutton et al., 2009). Thus, both experimental autoimmune encephalomyelitis (EAE) and imiquimod (IMQ)-induced psoriasis require the presence of functional γδT17 cells (Bekiaris et al., 2013; Sutton et al., 2009). Similarly, tumor models have shown that γδT17 cells can have either protective (Wu et al., 2014) or pathogenic (Coffelt et al., 2015) roles depending on the nature of the cancer. In humans, although a unique innate γδT17 cell population has not been yet characterized, many groups have identified IL-17-producing γδ T cells in association with various disease states (Cai et al., 2011; Wu et al., 2014).

Genetic murine studies have shown that γδT17 cells develop in the embryonic thymus in a step-wise fashion, initially involving escape of epithelial selection (Turchinovich and Hayday, 2011), followed by the upregulation of a number of transcription factors, such as RORγt, SOX13 and cMAF, that regulate lineage commitment, specification and functional maturation (Malhotra et al., 2013; Zuberbuehler et al., 2019). Although T cell receptor (TCR) signaling is necessary for γδT17 cell development (Wencker et al., 2014), experimental evidence based on hypomorphic CD3 mice and anti-CD3/TCR antibody administration suggested that only weak TCR signals are required (Munoz-Ruiz et al., 2016; Sumaria et al., 2017). In addition, a recent study showed that TCR signaling is not important for lineage specification but for transition into the early immature stage (Spidale et al., 2018). γδT17 cell generation is restricted to the embryonic and neonatal thymus (Haas et al., 2012) with the bone marrow displaying low capacity to produce these cells (Cai et al., 2014).

STAT transcription factors act downstream of cytokine and growth factor receptors to regulate a plethora of key biological processes, including lymphocyte development and function (Stark and Darnell, 2012). STAT5 is encoded by two genes, *Stat5a* and *Stat5b*, giving rise to two highly homologous proteins with largely overlapping functions in mediating transcription of target genes, although STAT5B has a more dominant role in lymphoid cells as well as in cancer progression (de Araujo ED, 2019; Villarino et al., 2016). Mice deficient in both STAT5A and STAT5B have increased perinatal mortality and lack or display a severe reduction in many lymphocytic populations, such as αβ T, γδ T, regulatory CD4^+^ T cells (Treg), natural killer (NK) T, NK and also B cells (Hoelbl et al., 2006; Yao et al., 2006; Yao et al., 2007). Although mice deficient in either STAT5A or STAT5B show hampered development and function of a few major lymphocyte subsets, their phenotype is milder than the combined loss of the two isoforms (Imada et al., 1998; Villarino et al., 2016; Villarino et al., 2017). However, there is currently no mechanistic data regarding their individual contribution in γδ T cell biology. One of the suggested mechanisms for the dependence of developing γδ T cell progenitors on STAT5 is its ability to induce TCRγ rearrangements, due to three highly interspecies conserved inverted repeat STAT5 consensus sites within the *TCRγ* locus (Wagatsuma et al., 2015).

In humans, *STAT5*-associated loss of function mutations are predominantly restricted to STAT5B and these culminate in growth failure, due to impaired growth hormone receptor signaling, as well as immunodeficiency, metabolic dysregulation and autoimmune disorders as a result of Treg deficiency (Cohen et al., 2006; Hwa, 2016; Kanai et al., 2012). In contrast, *STAT5B* gain of function (GOF) mutations strongly correlate with mature T cell neoplasms (Pham et al., 2018) and have also been found in patients with neutrophilia or eosinophilia (Cross et al., 2019; Ma et al., 2017). In particular, the recurrent N642H GOF missense mutation within the Src homology 2 (SH2) domain of STAT5B results in enhanced and prolonged tyrosine phosphorylation (pY) in response to low doses of cytokines or growth factors, and is associated with poorer patient prognosis and increased risk of relapse (Bandapalli et al., 2014; Pham et al., 2018; Rajala et al., 2013). Interestingly, *STAT5B* GOF mutations are relatively frequent in aggressive γδ T cell lymphoma subtypes, such as hepatosplenic T cell lymphoma (Nicolae et al., 2014), monomorphic epitheliotropic intestinal T cell lymphoma (Kucuk et al., 2015; Nairismagi et al., 2016) and primary cutaneous γδ T cell lymphoma (Kucuk et al., 2015). Notably, approximately 20% of identified N642H mutations occur in γδ T cell derived lymphomas (de Araujo ED, 2019).

Herein, we show that STAT5 is critically required for the progression and expansion of γδT17 cells through neonatal life in the intestine and periphery. We provide evidence that intestinal γδT17 cells upregulate Tbet upon entry into the lamina propria after birth and co-express the cytokines IL-17, IL-22 and IFNγ in a mechanism dependent on STAT3 and retinoic acid. Furthermore, loss of γδT17 cells due to STAT5 deficiency results in resistance to experimental autoimmune encephalomyelitis in adult mice. Importantly, we show that STAT5A promotes γδT17 cell expansion and downregulates intestinal Tbet favoring a type 17 program, whereas STAT5B favors IFNγ-producing γδ populations and increases intestinal Tbet expression. Collectively, our data suggest that neonatal life is a critical window of development and tissue specification for γδT17 cells, and that this process is tightly regulated by STAT5.

## Results

### STAT5 regulates the neonatal expansion of γδT17 cells

In order to test the importance of STAT5 in RORγt expressing γδ T cells, we crossed RORγt-Cre mice (Eberl and Littman, 2004) with mice floxed for both STAT5a and STAT5b (RORγt^CRE^-STAT5^F/F^) (Cui et al., 2004) and analyzed the numbers of LN and skin γδT17 cells. We found that compared to littermate controls (Cre^−^), RORγt^CRE^-STAT5^F/F^ mice (Cre^+^) contained severely reduced numbers of γδT17 cells defined phenotypically as CD27^−^CD44^+^ in the LN and CCR6^+^CD3^+^ in the skin (Fig. 1A-B). This was confirmed by the near complete lack of IL-17-expressing γδ T cells in the LN (Fig. 1C). Deficiency in STAT5 equally affected both Vγ4^+^ and Vγ4^−^ subsets of γδT17 cells (not shown). Interestingly, RORγt^CRE^-STAT5^F/F^ mice had a concomitant increase in IFNγ-expressing γδ T cells (Fig. 1C). In RORγt^CRE^-STAT5^F/F^ mice, deletion of STAT5 in CD4^+^ and CD8^+^ T cells was not complete (Fig. S1A). Insufficient deletion in the αβ T cell compartment using RORγt^CRE^ deleter mice has also been demonstrated by others (Guo et al., 2014). Consequently, we did not observe differences in the numbers of TCRβ^+^CD4^+^CCR6^+^ cells, which are enriched for T-helper-17 (Th17) cells (Fig S1B), or in the frequency of IFNγ-producing CD4^+^ T cells (Fig S1B). Surprisingly, and also in agreement with previous observations (Laurence et al., 2007), the percentage of IL-17A-producing CD4^+^ T cells was higher even when STAT5 was only partially deleted (Fig. S1B). Finally, to determine whether the defect we observed in RORγt^CRE^-STAT5^F/F^ mice was intrinsic to the γδT17 population we generated mixed bone marrow chimeras using CD45.1^+^ wild-type and Cre^+^ CD45.2^+^ donors and analyzed lymph nodes and skin 12 weeks later. We found that by comparison to wild-type, RORγt^CRE^-STAT5^F/F^ BM failed to generate γδT17 cells suggesting that the STAT5-associated defect is cell intrinsic (Fig. S2A). It is noteworthy that in the skin we could not detect any γδT17 cells originating from RORγt^CRE^-STAT5^F/F^ BM (Fig. S2A).

**Figure 1.**
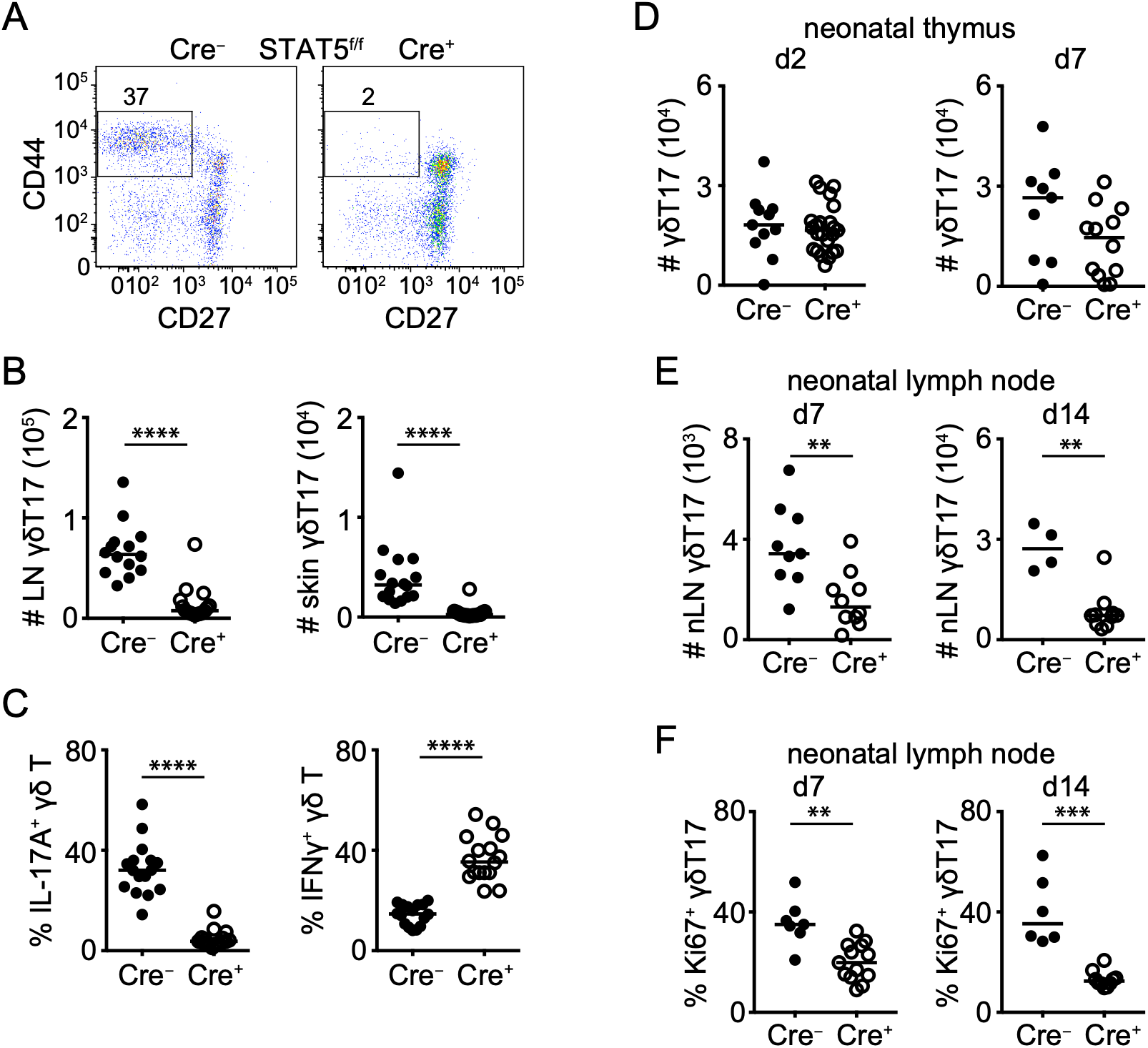
STAT5 is necessary for the neonatal expansion of γδT17 cells. Flow cytometric analysis of γδ T cells in RORγt^CRE^-STAT5^F/F^ (Cre^+^) and littermate control mice (Cre^−^). In graphs each symbol represents a mouse and line the median. **p < 0.01, ***p < 0.001, ****p < 0.0001 using Mann-Whitney test. (A) Expression of CD27 and CD44 in order to identify CD27^−^CD44^+^ γδT17 cells in the LN. Numbers indicate percent of CD27^−^CD44^+^ within the γδ T cell compartment. (B) Numbers of γδT17 cells in the LN (staining as in A) and skin. In the skin, γδT17 cells were identified as CD45^+^CD3^Low^Vγ5^−^TCRβ^−^TCRγδ^+^CCR6^+^. (C) Expression of IL-17A and IFNγ within the LN γδ T cell compartment. (D) Numbers of CD27^−^CD44^+^ γδT17 cells in day 2 and day 7 old thymi. (E) Numbers of CD27^−^CD44^+^ γδT17 cells in day 7 and day 14 old LN. (F) Frequency of Ki67^+^RORγt^+^ or Ki67^+^CCR6^+^ γδT17 cells within the CD44^+^TCRγδ^+^ compartment in day 7 and day 14 old LN.

We next investigated whether reduced γδT17 cell numbers in the absence of STAT5 were due to a developmental defect. We therefore examined newborn thymi from RORγt^CRE^-STAT5^F/F^ and littermate control mice and found no differences in γδT17 cellularity (Fig. 1D) or IL-17A expression (Fig. S2B). Expression of both *Stat5a* and *Stat5b* was significantly lower in RORγt^CRE^-STAT5^F/F^ γδT17 cells sorted from new born thymi compared to Cre^−^ controls or CD27^+^ γδ T cells (Fig. S2C). This suggested that the major impact of STAT5 occurs extrathymically. We thus analyzed neonatal mice and found a significant decrease in LN γδT17 cell numbers in 7 and 14 day old mice (Fig. 1E). Assessment of proliferation by Ki67 staining showed that γδT17 cells in RORγt^CRE^-STAT5^F/F^ neonatal mice displayed reduced turnover compared to controls (Fig. 1F). Furthermore, expression of the anti-apoptotic STAT5 target gene product BCL2 (Tripathi et al., 2010; Yao et al., 2006) was reduced in neonatal STAT5-deficient γδT17 cells (Fig. S2D), suggesting impaired survival. Collectively, we demonstrate that STAT5 is important for the turnover and survival of γδT17 cells during neonatal and adult life.

### Differential regulation of γδT17 and CD27^+^ γδ T cells by STAT5A and STAT5B

Since deficiency in STAT5 resulted in near complete loss of γδT17 cells, we next examined the influence of hyperactive STAT5 expression. We utilized two established models of STAT5 hyperactivity whereby the *Vav1* promoter drives the expression of (a) high or low copies of the hyperactive S710F STAT5A mutant (Maurer et al., 2019; Onishi et al., 1998), or (b) human wild-type (WT) or the hyperactive N642H STAT5B mutant (Pham et al., 2018). Constitutively high levels of hyperactive STAT5A resulted in very high numbers of γδT17 cells in LN, but hyperactivation of STAT5A had a considerably smaller impact on CD27^+^ γδ T cells (Fig. 2A-B). In contrast, constitutive expression of hyperactive STAT5B increased the numbers of CD27^+^ γδ T but had a smaller effect on γδT17 cells (Fig. 2A-B). When we analyzed cytokine expression we found that IFNγ was only induced by hyperactive STAT5B (Fig. S2E), whereas IL-17A could be induced at high levels both by hyperactive STAT5A as well as WT STAT5B expression (Fig. 2C). However, hyperactive STAT5B did not induce IL-17A expression (Fig. 2C).

**Figure 2.**
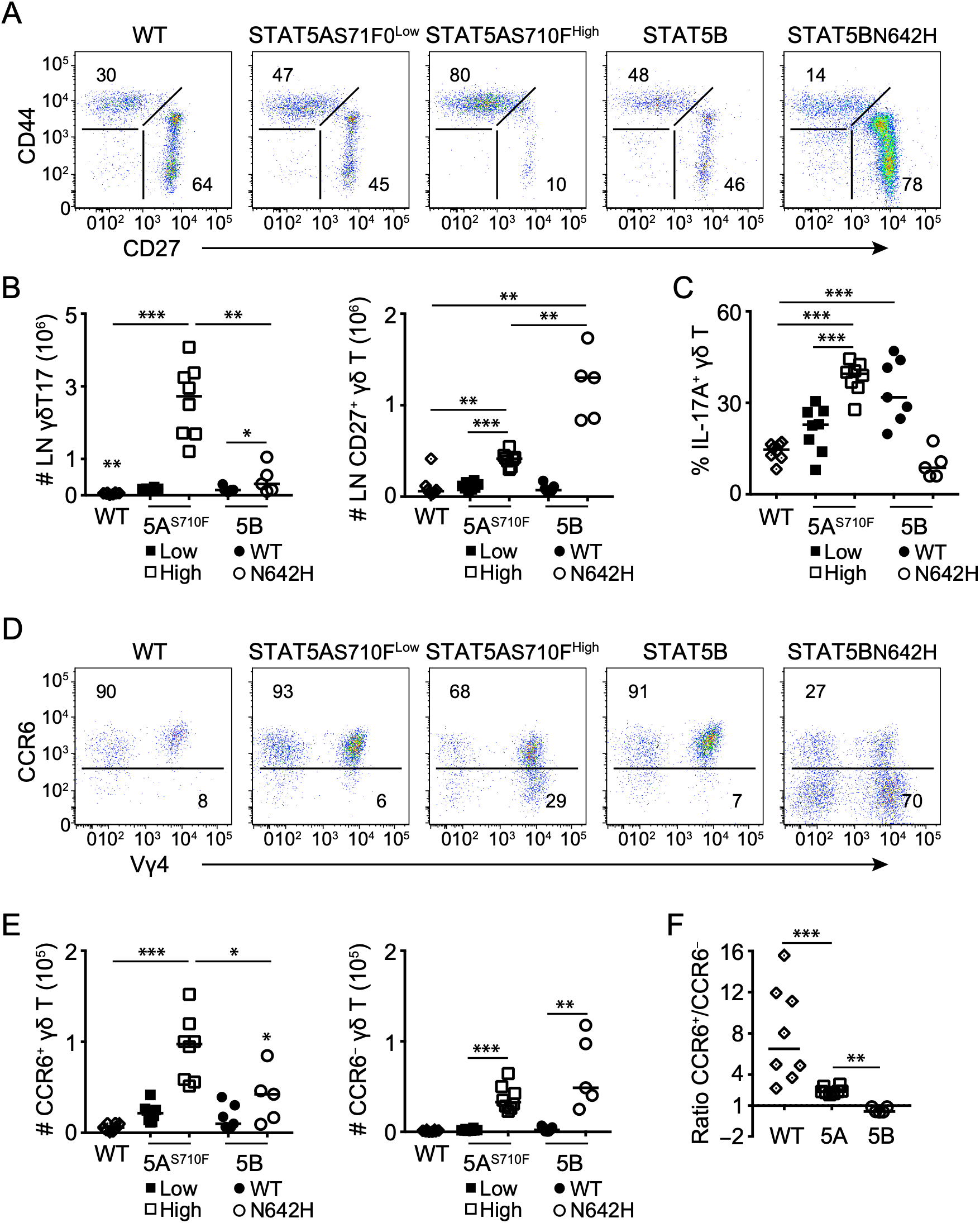
Differential impact of STAT5A and STAT5B on γδT17 and CD27^+^ γδ T cells. Flow cytometric analysis of γδ T cells in mice that are either wild-type (WT) or constitutively express under the *Vav1* promoter one of the following forms of STAT5: low (STAT5AS710F^Low^) or high (STAT5AS710F^High^) copy number of the hyperactive STAT5A S710F mutant, or human wild-type (STAT5B) or the hyperactive N642H STAT5B mutant (STAT5BN642H). In graphs, each symbol represents a mouse and line the median. *p < 0.05, **p < 0.01, ***p < 0.001 using Mann-Whitney test. (A) Expression of CD27 and CD44 in order to identify CD27^−^CD44^+^ γδT17 cells in the LN. Numbers indicate percent of CD27^−^CD44^+^ or CD27^+^CD44^−^ within the γδ T cell compartment. (B) Numbers of γδT17 (staining as in A) and CD27^+^ cells in the LN (** in WT denotes difference by comparison to STAT5BN642H). (C) Expression of IL-17A within the LN γδ T cell compartment. (D) Expression of CCR6 and Vγ4 in skin γδT17 cells (staining as in Fig. 1B). Numbers indicate percent of CCR6^+^ or CCR6^−^ cells within the γδ T cell compartment. (E) Numbers of CCR6^+^ and CCR6^−^ cells identified in D (* in STATBBN642H denotes significant difference by comparison to WT). (F) Ratio of CCR6^+^ over CCR6^−^ cells in WT compared to STAT5AS710F^High^ (5A) and STAT5BN642H (5B) mice.

Similar to the LN, skin γδT17 cell numbers were greatly enhanced by hyperactive STAT5A whereas STAT5B had a milder impact (Fig. 2D-E). With the exception of Vγ5^+^ dendritic epidermal T cells, CCR6^+^ γδT17 cells are the only γδ population in the skin. However, mice expressing hyperactive STAT5B, and to a lesser extent mice expressing hyperactive STAT5A, contained CCR6^−^ γδ T cells that were either Vγ4^+^ or Vγ4^−^ (Fig. 2D-F). Collectively, our data pinpoint towards a dominant role of STAT5A in supporting γδT17 cells in LN and skin, with STAT5B supporting mainly IFNγ-producing and CCR6^−^ γδ T cells.

### RORγt^CRE^-STAT5^F/F^ mice are resistant to experimental autoimmune encephalomyelitis

γδT17 cells have been implicated in the pathogenesis of experimental autoimmune encephalomyelitis (EAE) (Petermann et al., 2010; Sutton et al., 2009) and we therefore investigated how well RORγt^CRE^-STAT5^F/F^ mice responded to MOG (myelin oligodendrocyte glycoprotein)-induced EAE. We found that compared to littermate controls, RORγt^CRE^-STAT5^F/F^ mice were resistant to EAE symptoms (Fig. 3A). This correlated with significantly reduced γδT17 cells in the LN and brain at days 11 and 21 after immunization (Fig. 3B-D). As expected, γδ T cell associated IL-17A production was significantly reduced at all time points in mice lacking STAT5 (Fig. 3E). Although it has been recently suggested that inflammatory conditions during EAE can *de novo* regenerate γδT17 cells (Papotto et al., 2017a), our data suggest that in the absence of STAT5, γδT17 cell regeneration cannot occur.

**Figure 3.**
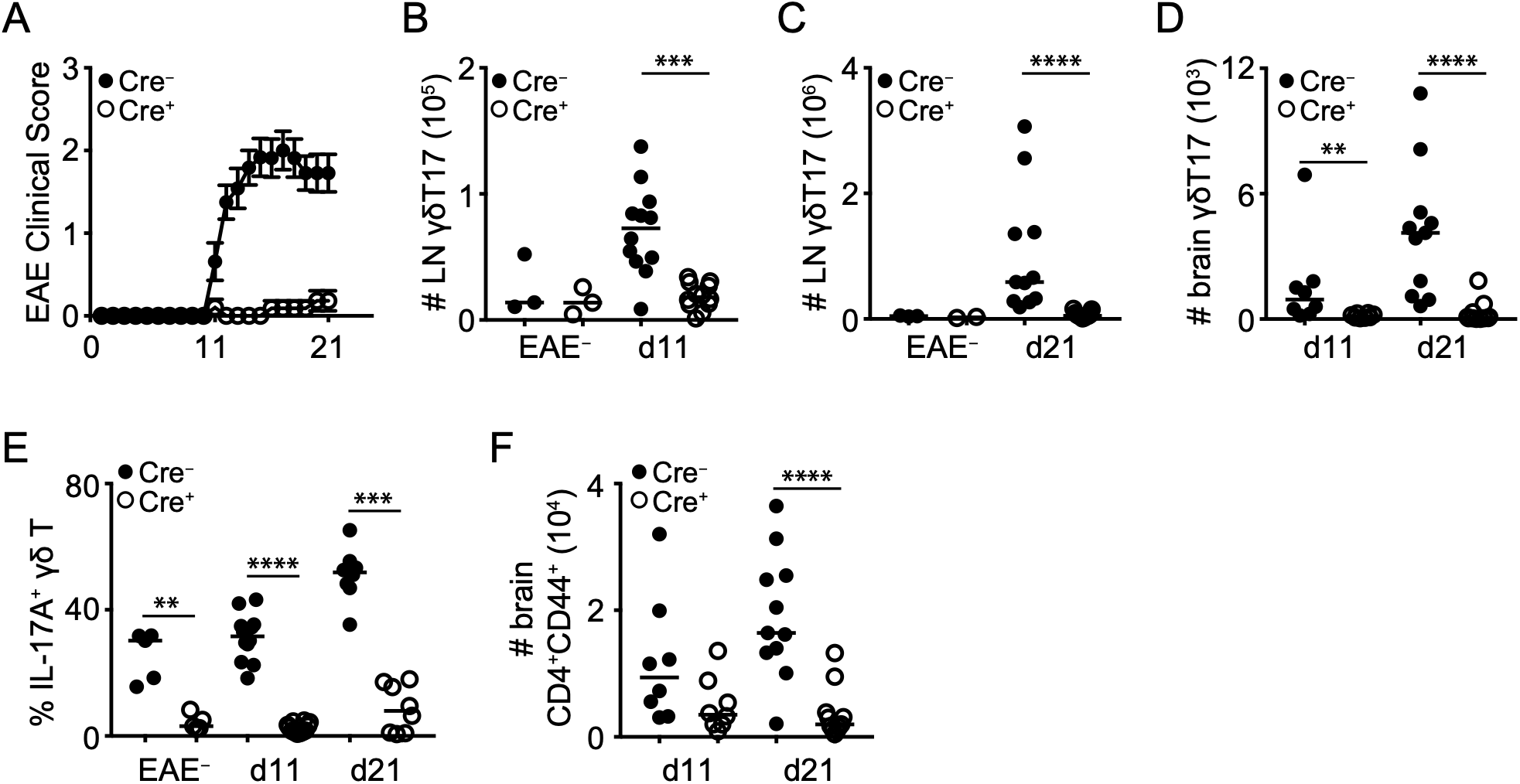
RORγt^CRE^-STAT5^F/F^ mice are resistant to EAE. Disease progression and flow cytometric analysis of γδ and CD4^+^ T cells in RORγt^CRE^-STAT5^F/F^ (Cre^+^) and littermate control mice (Cre^−^) that had been previously immunized with 50 µg MOG peptide in CFA and 200 ng Pertussis toxin. In graphs, each symbol represents a mouse and line the median (except in A). **p < 0.01, ***p < 0.001, ****p < 0.0001 using Mann-Whitney test. (A) Clinical symptoms of EAE until day 21 post immunization. Data is pool of 20 mice per genotype and shown as mean±sem. Statistical analysis was performed using 2-way ANOVA with Bonferroni’s multiple comparisons test. ANOVA p-value < 0.0001; Bonferroni’s test returned significance for days 11-21 with day 11 p = 0.003 and days 12-21 p < 0.0001. (B-C) Numbers of γδT17 cells in the LN (staining as in Fig. 1A) of unimmunized controls (EAE^−^) and at 11 (B) and 21 (C) days after immunization. (D) Numbers of γδT17 cells in the brain (identified as CD45^+^CD3^+^TCRβ^−^TCRγδ^+^CD44^+^) at days 11 and 21 after immunization. (E) Expression of IL-17A within the LN γδ T cell compartment of unimmunized controls (EAE^−^) and at 11 and 21 days after immunization. (F) Numbers of CD4^+^CD44^+^CD3^+^TCRβ^+^ cells in the brain at days 11 and 21 after immunization.

It has been shown that in addition to their direct contribution to EAE pathogenesis, γδT17 cells are required for optimal Th17 responses (Sutton et al., 2009). We therefore interrogated the CD4^+^ T cell response in the LN and brain of RORγt^CRE^-STAT5^F/F^ and littermate control mice during EAE. We found that the numbers and cytokine production of CD4^+^ T cells were not affected in the LN (Fig. S1C-D), which may be a reflection of the levels of STAT5 still detectable in these cells (Fig. S1A). As expected from the clinical score and the lack of pro-inflammatory γδT17 cells, there was a profound reduction in CD4^+^ T cell numbers within the brain of RORγt^CRE^-STAT5^F/F^ mice (Fig. 3F). Collectively, our data show that loss of γδT17 cells due to STAT5 deficiency is associated with dramatically reduced inflammatory responses in the EAE model.

### Intestinal lamina propria γδT17 cells express Tbet and require STAT3 and retinoic acid for cytokine production

Besides the skin and peripheral lymphoid tissues, γδ T cells with type 3 functionality have been described in the mucosa such as the lung and gut (Sheridan et al., 2013; Sutherland et al., 2014). We therefore wanted to test whether STAT5 regulated γδT17 cells specifically in the intestinal lamina propria (LP). In order to avoid potential differences in γδT17 surface markers in the gut, we stained small intestinal and colonic LP (sLP and cLP respectively) for RORγt and Tbet and compared this to peripheral LNs. Surprisingly, we found that many RORγt^+^ γδ T cells in the gut co-expressed Tbet (Fig. 4A). We additionally confirmed the presence of RORγt^+^Tbet^+^ γδ T cells by generating double transgenic mice reporting GFP and AmCyan under control of the promoters for RORγt and Tbet, respectively (Fig S3A). By transcription factor staining analysis we found that the RORγt^+^Tbet^+^ γδ T cell population was more prevalent in the ileum and proximal colon (Fig. 4B), which contrasted with the distribution of RORγt^−^Tbet^+^ γδ T cells in the same locations (Fig. S3B). In order to investigate which factors regulate the expression of Tbet, we analyzed mice deficient in Toll-like receptor and IL-12 signaling as well as mice depleted of their intestinal microbial flora. We found that expression of Tbet was independent of the microbiota (Fig. S3C), MyD88, TRIF and IL-12 signaling (Fig. S3D).

**Figure 4.**
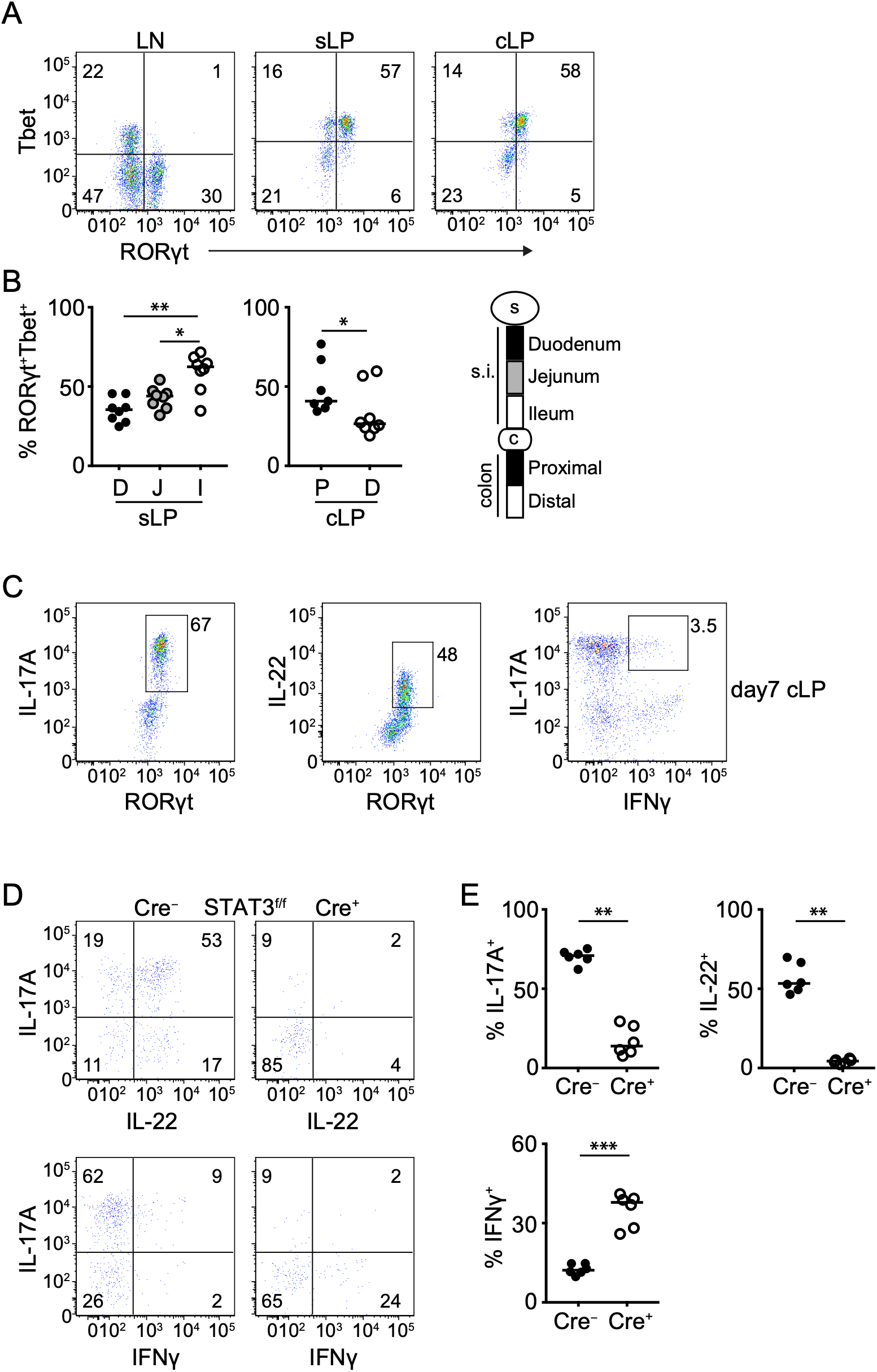
Intestinal γδT17 cells express Tbet and require STAT3 for IL-17A and IL-22. Flow cytometric analysis of LN and intestinal γδ T cells in WT or in RORγt^CRE^-STAT3^F/F^ (Cre^+^) and littermate control mice (Cre^−^). In graphs, each symbol represents a mouse and line the median. Cytokine detection was performed following IL-23 re-stimulation. *p < 0.05, **p < 0.01, ***p < 0.001, ****p < 0.0001 using Mann-Whitney test. (A) Expression of RORγt and Tbet within the γδ T cell compartment of the LN, sLP and cLP. Numbers indicate percent of RORγt and Tbet expression. (B) Frequency of RORγt^+^Tbet^+^ cells within the γδ T cell compartment in the indicated small intestinal and colonic segments. (C) Expression of RORγt and IL-17A, RORγt and IL-22, or IL-17A and IFNγ in the γδ T cell compartment of cLP from day 7 old mice (representative of two experiments). (D) Expression of IL-17A and IL-22 (top) or IL-17A and IFNγ (bottom) in RORγt^CRE^-STAT3^F/F^ (Cre^+^) and littermate control mice (Cre^−^). Numbers indicate percent of positive expression. (E) Frequency of IL-17A^+^, IL-22^+^ and IFNγ^+^ γδ T cells in RORγt^CRE^-STAT3^F/F^ (Cre^+^) and littermate control mice (Cre^−^).

In agreement with their innate nature, intestinal γδT17 cells produced IL-17A, IL-22 and IFNγ as early as 7 days after birth (Fig. 4C), indicating a functional γδT17 population that has acquired the ability to produce IFNγ at steady-state. Using RORγt^CRE^-STAT3^F/F^ mice, we showed that production of both IL-17A and IL-22 was STAT3-dependent (Fig. 4D-E and S4A-B). One of the key factors regulating IFNγ-expressing cells is retinoic acid (RA) (Brown et al., 2015). We therefore interrogated mice possessing a RA receptor (RAR) dominant negative (RARdn) transgene, which prevents active RARα signaling (Rajaii et al., 2008) in RORγt-expressing cells (RORγt^CRE^-RARdn^F/F^ mice). We found that loss of RA signaling was associated with reduced overall expression of IFNγ (Fig. 5A and S4C) as well as reduced frequency of IL-17A^+^IFNγ^+^ and IL-22^+^IFNγ^+^ cells (Fig. 5B and S4D). In contrast, deficiency in RA signaling resulted in significantly increased frequency of IL-17A^+^IFNγ^−^ and IL-22^+^IFNγ^−^ Tbet^+^ γδT17 cells in the colon and small intestine (Fig. 5B and S4D). Collectively, this data indicates that lamina propria Tbet^+^ γδT17 cells are innate cells that can co-produce IL-17, IL-22 and IFNγ, and that their cytokine expression profile is regulated by STAT3 and RA.

**Figure 5.**
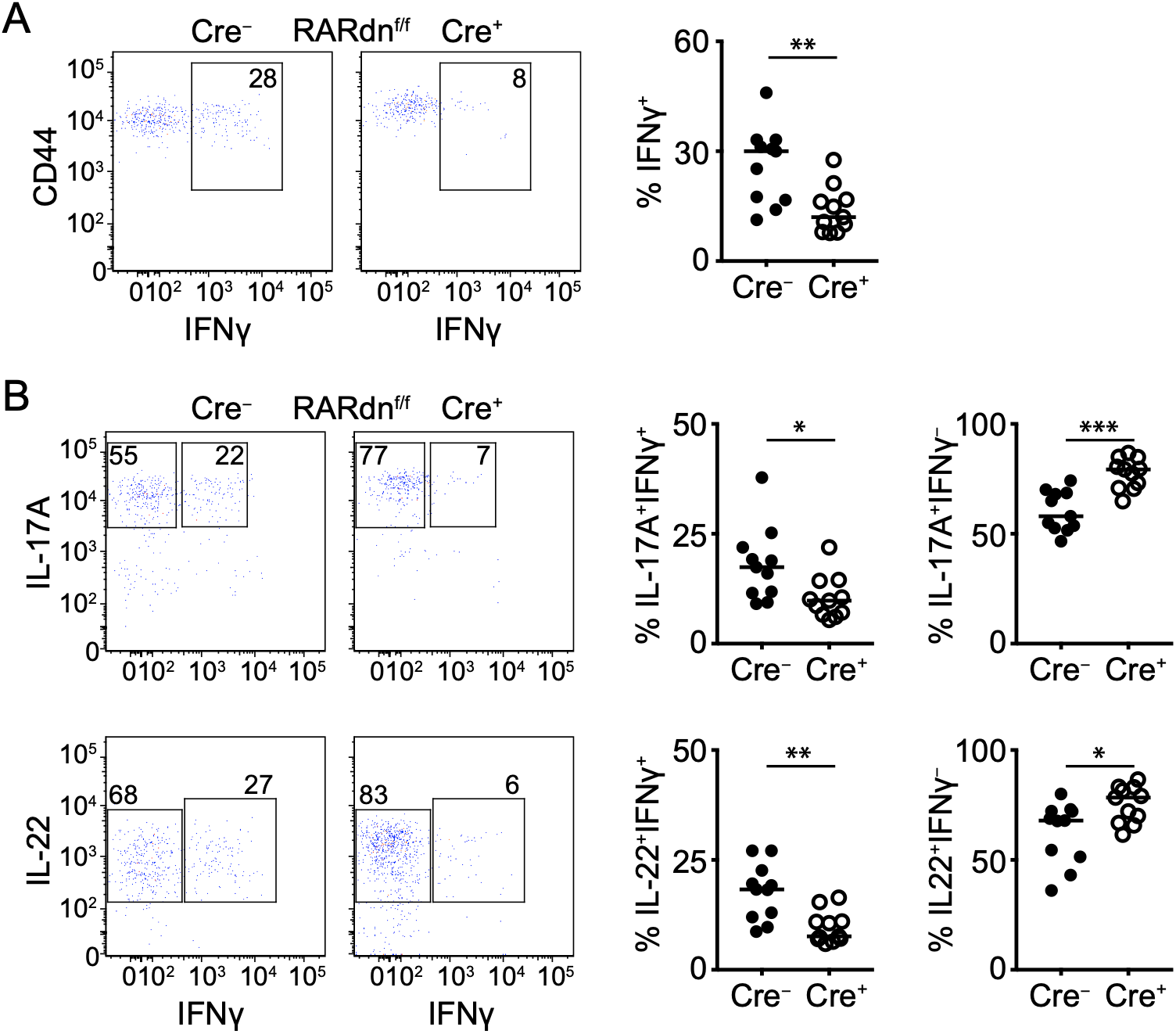
Retinoic acid receptor signaling regulates IFNγ production in intestinal Tbet^+^ γδT17 cells. Flow cytometric analysis of cytokine expression in colonic γδ T cells in RORγt^CRE^-RARdn^F/F^ (Cre^+^) and littermate control mice (Cre^−^) following IL-23 stimulation. In graphs, each symbol represents a mouse and line the median. *p < 0.05, **p < 0.01, ***p < 0.001 using Mann-Whitney test. (A) Expression of CD44 and IFNγ (dot plots) and frequency of IFNγ ^+^ γδ T cells (graph) in RORγt^CRE^-RARdn^F/F^ (Cre^+^) and littermate control mice (Cre^−^). (B) Expression of IL-17A and IFNγ (top dot plots) or IL-22 and IFNγ (bottom dot plots) with graphical representation of the frequency of IL-17A^+^IFNγ^+^ and IL-17A^+^IFNγ^−^ or IL-22^+^IFNγ^+^ andIL-22^+^IFNγ^−^ γδ T cells in RORγt^CRE^-RARdn^F/F^ (Cre^+^) and littermate control mice (Cre^−^).

### STAT5 regulates Tbet expression and determines the progression of intestinal γδT17 cells through neonatal development

Following the identification of a distinct gut-specific γδT17 population, we aimed to understand their dependence on STAT5. Similar to LNs, RORγt-expressing γδ T cells, irrespective of Tbet, were drastically and significantly reduced from the sLP and cLP of RORγt^CRE^-STAT5^F/F^ mice (Fig. 6A-B). Analysis of GOF STAT5A and STAT5B mice revealed that hyperactive STAT5A downregulated Tbet in RORγt^+^ cLP γδ T cells, whereas hyperactive STAT5B enhanced it (Fig. 6C) suggesting a YIN/YANG regulation in RORγt^+^ cLP γδ T cells by STAT5A versus STAT5B. Hyperactive STAT5A preferentially expanded RORγt^+^ cells in the gut whereas hyperactive STAT5B favored Tbet-expressing γδ T cells irrespective of whether they expressed RORγt or not (Fig. 6D-F).

**Figure 6.**
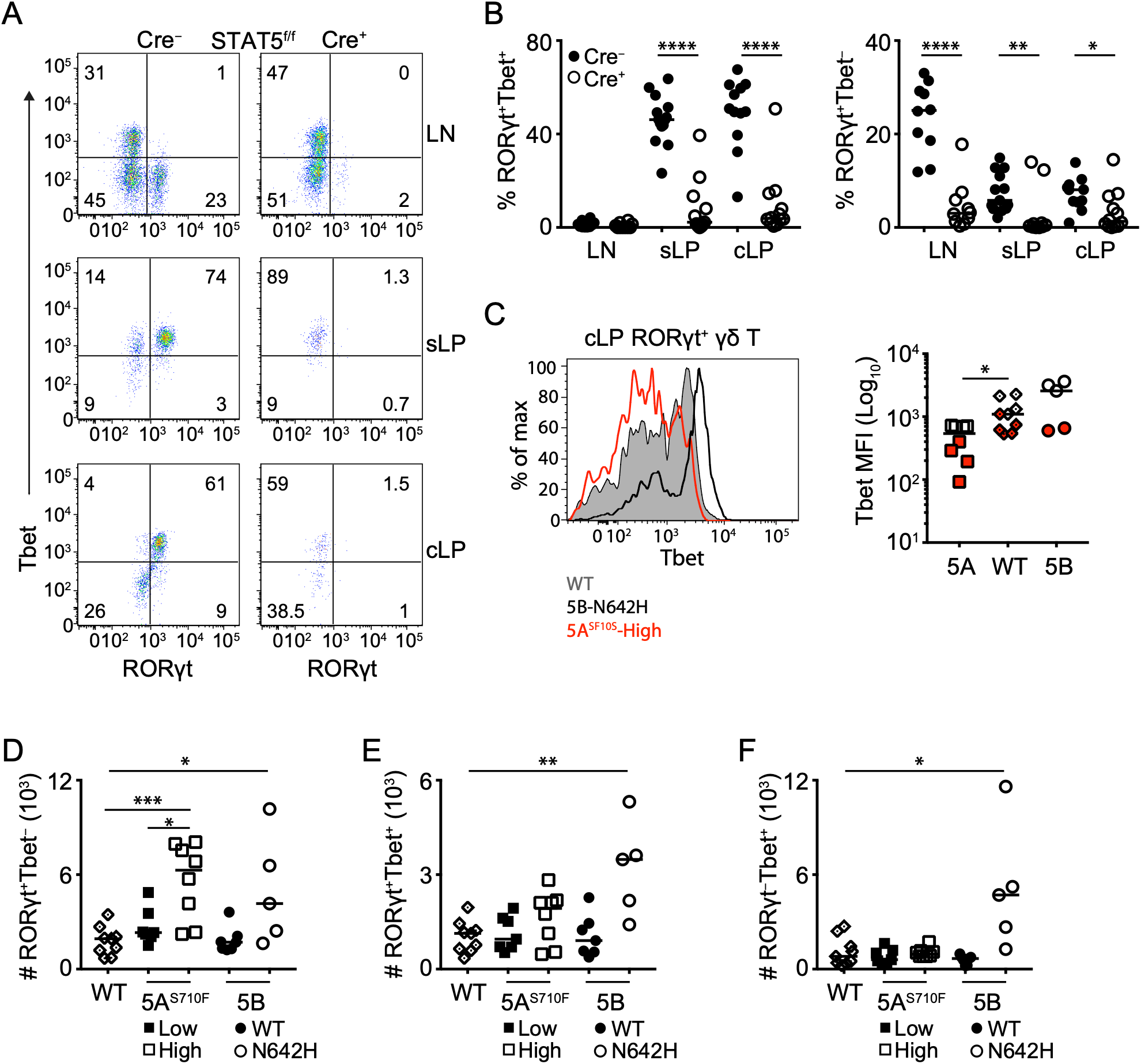
STAT5 is a critical determinant of Tbet-expressing intestinal γδT17. Flow cytometric analysis of intestinal γδ T cells in RORγt^CRE^-STAT5^F/F^ (Cre^+^) and littermate control mice (Cre^−^) in STAT5A and STAT5B hyperactive mutant mice as described in Figure 2. In graphs, each symbol represents a mouse and line the median. *p < 0.05, **p < 0.01, ***p < 0.001, ****p < 0.0001 using Mann-Whitney test. (A) Expression of RORγt and Tbet within the γδ T cell compartment of the LN, sLP and cLP. Numbers indicate percent of RORγt and Tbet expression. (B) Frequency of RORγt^+^Tbet^+^ and RORγt^+^Tbet^−^ cells within the γδ T cell compartment in LN, sLP and cLP. (C) Expression of Tbet (histogram) and Tbet mean fluorescent intensity (MFI) (graph) in RORγt^+^ cLP γδ T cells from WT, STAT5AS710F^High^ (5a) or STAT5BN642H (5b) mice as described in Figure 2. In graph, colors indicate two different experiments. (D) Numbers of RORγt^+^ (E), RORγt^+^Tbet^+^ (F) and RORγt^−^Tbet^+^ γδ T cells in the cLP of the indicated STAT5A and STAT5B hyperactive mutant mice or WT control mice.

Next, we sought to determine whether STAT5 also regulated RORγt^+^Tbet^+^ γδ T cells neonatally. We therefore analyzed neonatal gut at different time points and found that 1-2 days after birth γδ T cells in the colon and small intestine expressed either RORγt or Tbet but not both (Fig. 7A-C and S5A-C). Tbet was induced in RORγt-expressing cells at day 4 and stabilized to adult levels within the first week of life (Fig. 7A-B and S5A-B). Expression of Tbet at neonatal day 4 coincided with a rapid increase in cell proliferation, which was blunted in the absence of STAT5 (Fig. 7D and S5D). RORγt^CRE^-STAT5^F/F^ mice did not upregulate Tbet and failed to sustain a RORγt^+^ γδ T cell population after birth (Fig. 7A-C and S5A-C). However, despite their functional presence in the neonatal gut, RORγt-expressing γδ T cells were not necessary for protection against early life infection with the attaching and effacing bacterium *Citrobacter rodentium* (Fig. S6A-C).

**Figure 7.**
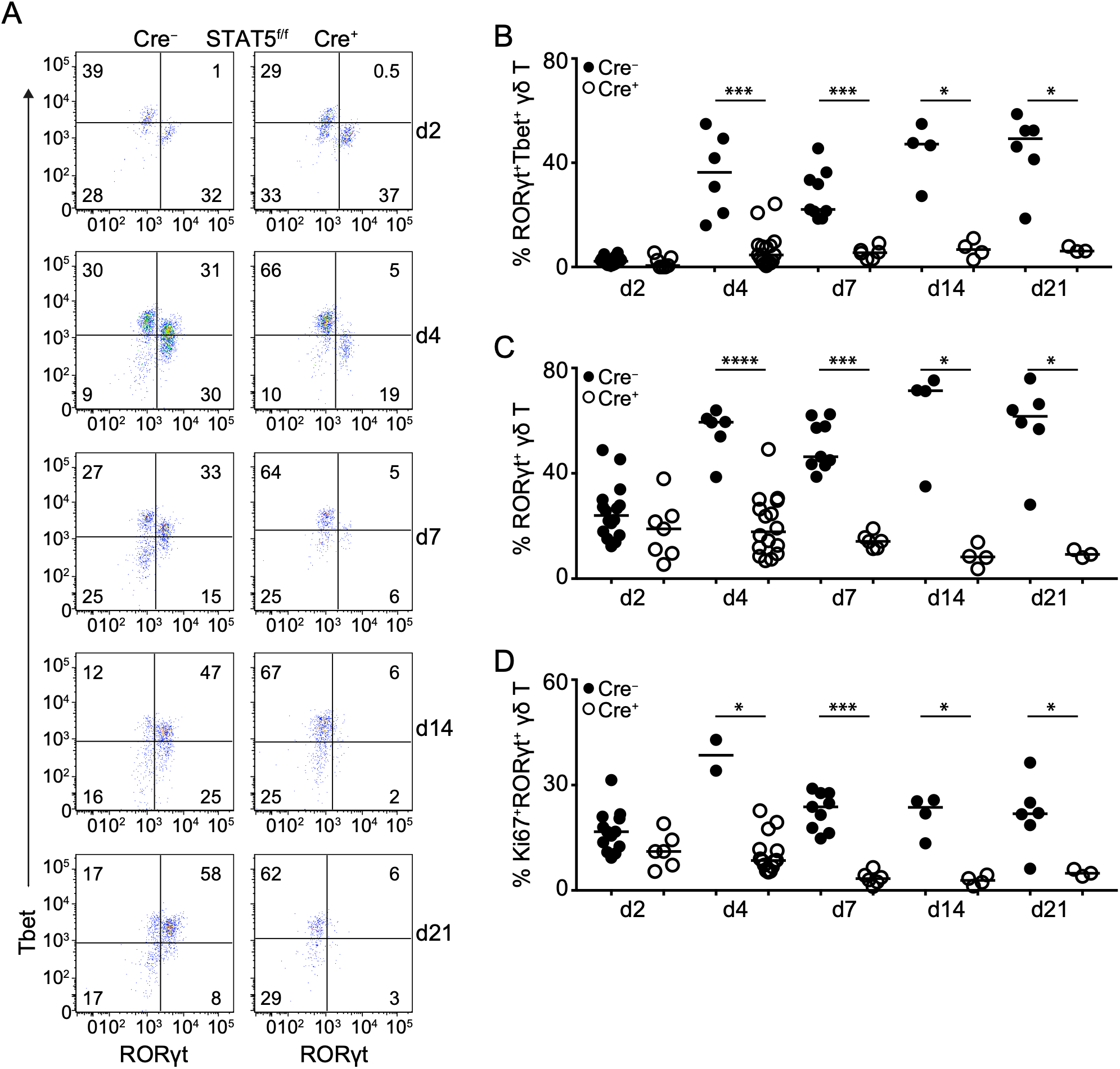
STAT5 regulates the neonatal fate of intestinal Tbet^+^ γδT17 cells. Flow cytometric analysis of colonic γδ T cells in RORγt^CRE^-STAT5^F/F^ (Cre^+^) and littermate control mice (Cre^−^) during neonatal ontogeny. Day of birth is counted as day(d)1. In graphs each symbol represents a mouse and line the median. *p < 0.05, **p < 0.01, ***p < 0.001, ****p < 0.0001 using Mann-Whitney test. (A) Expression of RORγt and Tbet within the γδ T cell compartment of cLP at the indicated days after birth. Numbers indicate percent of RORγt and Tbet expression. (B) Frequency of cLP RORγt^+^Tbet^+^ γδ T cells at the indicated days after birth. (C) Frequency of cLP RORγt^+^ γδ T cells (including Tbet^+^) at the indicated days after birth. (D) Frequency of cLP Ki67^+^RORγt^+^ γδ T cells at the indicated days after birth.

Collectively, our data demonstrate that during neonatal life STAT5 acts as a molecular checkpoint to promote proliferation of intestinal γδT17 cells. Moreover, our data reveal an interesting balance between STAT5A and STAT5B, which appear to have opposing roles in the regulation of Tbet expression, thereby differentially coordinating tissue specificity of γδT17 cells.

## Discussion

In the present study we demonstrate that STAT5 is a critical regulator of IL-17-producing γδ T cells in the periphery, skin and gut. STAT5 was necessary during neonatal life in order to sustain proliferation and survival of γδT17 cells. Transgenic reconstitution of hyperactive STAT5 variants showed that STAT5A preferentially sustains γδT17 whereas STAT5B promotes IFNγ-producing γδ T cells. Physiologically, hampered development of γδT17 cells due to STAT5 loss resulted in near complete resistance to EAE pathology and prevented Th17 cells from infiltrating the brain. Furthermore, we discovered that intestinal lamina propria γδT17 cells co-express the type 1 transcription factor Tbet and can produce IL-17, IL-22 and IFNγ in a STAT3- and RA-dependent mechanism. Intestinal γδT17 cells upregulate Tbet during the first week of life and are strictly STAT5-dependent for their neonatal development. Furthermore, expression of Tbet is under the antagonistic control of STAT5A and STAT5B.

STAT5 is a major signaling component downstream of many cytokine and growth factor receptors and is therefore involved in the development of lymphocyte lineages (Rani and Murphy, 2016). Hence, both mice and humans with STAT5-associated deficiencies are severely immunocompromized (Imada et al., 1998; Kofoed et al., 2003). T cells of the γδ lineage are reduced in the thymus and lymphoid tissues of full STAT5-deficient mice (Hoelbl et al., 2006), and this has been attributed to a failure to successfully rearrange the TCR early during embryonic development (Wagatsuma et al., 2015). However, our data show that γδT17 cells require STAT5 signaling to expand and survive after they exit the thymus. This suggests that γδ T cell subsets rely on STAT5 during different steps of their development and differentiation, presumably reflecting cytokine niches within the local microenvironment.

Detailed molecular and phenotypic studies utilizing mice deficient in either STAT5A or STAT5B have shown that despite their many commonalties, particularly at the genome-wide level, the two STAT5 gene products can display cell-specific functions (Villarino et al., 2016; Villarino et al., 2017). Thus, in CD4^+^ T cells STAT5B has a dominant role in orchestrating differentiation and function. In this regard, our data show that both STAT5A and STAT5B can have dominant and differential roles in γδ T cells depending on the specific subset. Whereas STAT5A regulated almost exclusively γδT17 cells and downregulated intestinal Tbet expression, STAT5B had a prevailing effect on IFNγ-expressing γδ populations. This suggests that unlike in CD4^+^ helper and innate lymphoid cells, STAT5A and STAT5B display significant, differential regulatory functions in γδ T cell subsets, and further pinpoints to the distinct molecular, functional and developmental requirements of γδT17 compared to non-IL-17-producing subsets. The unique roles that we uncovered herein for STAT5A and STAT5B suggest that they display cell-specific functions and can have context-dependent, non-redundant roles in generating robust immune responses. The genetic and cellular tools that we used herein will be crucial to illuminate the specific biological functions of these two highly species-conserved proteins that play indispensable roles in infection, cancer and autoimmunity.

Although γδT17 cells develop in the embryonic thymus, previous reports suggested that they populate the skin and LNs after birth (Cai et al., 2014). Findings herein indicate that the neonatal period is critical for γδT17 cells to populate lymphoid and non-lymphoid tissues. A critical time window of opportunity has been suggested to exist during neonatal life when, upon exposure to microbiota, the immune system matures and develops cellular and humoral immunity (Al Nabhani et al., 2019; Torow and Hornef, 2017). The upregulation of Tbet in murine intestinal γδT17 cells within days after birth and its independence on the microbiota and TLR signals suggests alternative neonatal factors such as lactation, which is predominantly STAT5A-controlled (Haricharan and Li, 2014). Neonatal-specific cytokine milieus that activate STAT5 may also regulate Tbet expression in the developing gut. In this regard, hyperactive STAT5A downregulated Tbet whereas high STAT5B activity induced it, although whether this was direct or indirect through regulation of cell fate transcriptional regulators remains to be studied. Nevertheless, the identification of Tbet-expressing γδT17 cells at steady-state indicates a form of plasticity within this lineage that is regulated post-thymically and in a tissue-specific manner. The importance of Tbet in γδT17 cells is currently unknown. However, the co-expression of IFNγ suggests that acquisition of type 1 transcriptional and functional traits may give an advantage over infection, similar to innate lymphoid cells and Th1-transitioning Th17 cells.

Animal models have linked γδT17 cells to immune responses during inflammation, infection and cancer where they can be either protective or pathogenic (Papotto et al., 2017b). In the imiquimod model of psoriasis, γδT17 cells are important to drive disease; however, their pathogenic role can be redundant and compensated by other inflammatory cells (Sandrock et al., 2018). In the EAE mouse model, γδT17 cells have also been shown to contribute to pathogenicity (Petermann et al., 2010; Sutton et al., 2009). Herein, we provide evidence that γδT17 cells are necessary and non-redundant for full development of EAE symptoms and for the mobilization of Th17 cells to the brain. Despite their absence from all major organs from neonatal life onwards, other innate inflammatory cells could not compensate for their absence. γδ T cells have co-evolved alongside αβ T and B cells (Hirano et al., 2013), however their function has diversified and was imprinted by the specific tissue cues present in different locations. It is thus not surprising that the immunological response of different γδ T cell subsets will vary and will be essential or redundant depending on the inflammatory or infectious context.

In summary, we provide evidence that the neonatal microenvironment, acting in synergy with tissue-specific and STAT5-driven molecular cues, regulates the development, functional maturation and immunological importance of γδT17 cells.

## Acknowledgments

We thank Dr John O’Shea (NIH/NIAMS, MD) and Dr Stephen Shoenberger (La Jolla Institute for Immunology, CA) for critically reading the manuscript. This work, V.B. and DK were supported by the Lundbeck Foundation grant R163-2013-15201. D.K. was additionally supported by the Leo Foundation grant LF16020. J.R. and R.A. were supported by DTU PhD scholarships. R.M. and H.N. were supported by a private cancer metabolism grant donation from Liechtenstein and R.M., T.S. and B.M. were further supported by the Austrian Science Fund (FWF), grants SFB F4707 and SFB-F06105, Austria.

## Author Contributions

V.B. designed the study, oversaw all work, performed experiments and wrote the manuscript. D.K. designed and performed experiments, analyzed data and helped write the manuscript. J.R. and R.A. designed and performed experiments, analyzed data and helped write the manuscript. H.N., T.S., B.M. and R.M. generated the STAT5 GOF mutant strains, performed experiments, analyzed data and helped write the manuscript. E.C. and A.T. performed and analyzed experiments.

## Methods

### Mice

All animal breeding and experiments were performed in house at DTU and only after approval from the Danish Animal Experiments Inspectorate. RORγt^CRE^ and RORγt^GFP^ mice (Lochner et al., 2008) were provided by Professor Gerard Eberl and subsequently bred in house. STAT5^F/F^ mice were generated as previously described (Cui et al., 2004) and bred in house. STAT3^F/F^ mice were purchased from The Jackson Laboratory and bred in house. RARdn^F/F^ mice were generated as previously described (Rajaii et al., 2008), were provided by Professor William W. Agace, (Lund University, Sweden) and bred in house. Intestinal tissue from IL12^−/−^ mice (JAX stock #002692) was provided by Professor William W. Agace, (Lund University, Sweden). GOF STAT5a and STAT5b mice were generated as previously described (Maurer et al., 2019; Pham et al., 2018) and were bred at the University of Veterinary Medicine Vienna (Vienna, Austria). Frozen sperm samples from Tbet-AmCyan mice (Yu et al., 2015) were provided by Professor Jinfang Zhu (NIH/NIAID, MD) and after re-derivation mice bred in house. Intestinal tissue from mice deficient in TRIF and MyD88 was provided by Dr Katharina Lahl (DTU, Denmark).

### Cell culture media and buffers

All cell culture and single cell suspensions were prepared using RPMI 1640 (Invitrogen) supplemented with 10% heat inactivated fetal bovine serum (FBS)(Gibco), 20mM Hepes pH 7.4 (Gibco), 50 µM 2-mercaptoethanol, 2 mM L-glutamine (Gibco) and 10,000 U/ mL penicillin-streptomycin (Gibco). 10x Hank’s balanced salt solution (HBSS)(Gibco) was diluted to 1x with sterile nuclease-free water and supplemented with 15 mM HEPES pH 7.4 (Gibco) to prepare HBSS-HEPES while it was supplemented with 2mM EDTA, 15mM HEPES, 50µg/mL Gentamycin and 2% FBS to prepare HBSS-EDTA. Isotonic Percoll was prepared by mixing 90%v/v of Percoll (GE healthcare) with 9%v/v 10x HBSS and 1%v/v 1M HEPES pH 7.4. Isotonic Percoll was subsequently diluted with HBSS-EDTA to the desired concentration. FACS buffer was prepared by mixing 3% heat inactivated FBS with DPBS (Gibco).

### Isolation of lymphocytes from lymph nodes (LNs), thymus, skin, small intestine and colon

LNs were dissected, cleared off fat and crushed against a 70µm cell strainer to prepare single cell suspensions. Cell suspensions were then washed and filtered through a 40µm cell strainer. Cells were counted and 2.5×10^6^ cells were used for staining of surface antigens and flow cytometry analysis.

Thymus lobes from 1-day old pups were dissected and dissociated in supplemented RPMI using dissection microscope to prepare single cell suspension. The cell suspensions were filtered through a 40µm cell strainer and stained for surface antigens under sterile conditions before FACS sorting.

Skin lymphocytes were prepared from ears as follows: first, the dorsal and ventral sides of the ears were mechanically separated, they were subsequently cut into small pieces followed by enzymatic digestion with 0.25mg/ml collagenase IV, 0.166mg/ml hyaluronidase and 0.1mg/ml DNase I (all enzymes from Sigma-Aldrich) in supplemented RPMI for 1 hour at 37°C with constant stirring at 700 rpm. Undigested tissue was crushed against a 70µm cell strainer to prepare a single cell suspension. After washing, the cell pellet was re-suspended and filtered through a 40µm cell strainer to remove tissue debris and used for flow cytometry staining.

Small intestines and colons were dissected from adult mice and were HBSS-HEPES to remove intestinal contents. Fat and Peyer’s patches were removed before and then the tissues were open longitudinally and cut into small pieces of approximately 2-3 cm. Chopped tissue was washed 4 times (2 alternate cycles of 10 and 15 min each) using 15 mL of HBSS-EDTA buffer at 37°C in a shaking incubator. Tissue pieces were then digested using 0.3mg Liberase TM (Roche) and 0.15 mg of DNase (Sigma Aldrich) per preparation in 5mL supplemented RPMI for 40 minutes on the magnetic stirrer at 37°C. The resulting cell suspensions were filtered through 70µm cell strainers, collected in complete RPMI and subsequently pelleted by centrifugation. The cell pellets were then re-suspended in 5 mL 40% Percoll, layered on 4 mL of 70% Percoll, and centrifuged at 20 °C and 800 × g for 20 min with deceleration set to 0. Cells from the interphase were collected, washed once and then re-suspended in supplemented RPMI. For neonatal gut samples, cell suspensions, following digestions, were filtered through 70µm cell strainers and were then used directly.

### Experimental Autoimmune Encephalomyelitis

EAE was induced by sub-cutaneous injection of 50µg of MOG35-55 peptide in CFA, while 2 ng pertussis toxin were intra-peritoneally (i.p.) injected on the day of immunization and 2 days later. From day 11 after immunization and until day 21, mice were weighed and scored for clinical signs as follows: 0: no symptoms; 1: tail paralysis; 1.5: impaired righting reflex; 2: paralysis of one hind limb; 2.5: paralysis of both hind limbs; 3: paralysis of one fore limb; 3.5: paralysis of one fore limb and weak second for limb; 4: total limb paralysis.

Mice were euthanized at days 11 or 21 after immunization mice and were perfused with PBS. LN cells were isolated as described above. Brain tissue was mechanically minced and passed through a 70µm cell strainer to obtain a single cell suspension. Lymphocytes were separated using density gradient centrifugation with 47% Percoll (GE Healthcare), layered on 4 mL of 70% Percoll, and centrifuged at 20 °C and 900 × *g* for 30 min with deceleration set to 0.

### *In vitro* stimulation of lymphocytes

For LN lymphocytes, 10^7^ cells were cultured for 3.5 hours in the presence of 50ng/ml PMA (Sigma), 750ng/ml ionomycin (Sigma) and 1µL /mL BD Golgistop (containing monensin). Single cell suspensions from intestinal lamina propria were stimulated with 40 ng/ml of IL-23 (R&D Systems) for 3 hours followed by 50ng/ml PMA, 750ng/ml ionomycin and 1µL /mL BD Golgistop for an additional 3 hours. After 6 hours, cells were harvested and washed with PBS and used for flow cytometry staining. FoxP3 transcription factor staining kit (eBiosciences)

### Flow Cytometry

Cells were harvested by centrifugation at 400 g for 5 minutes at 4°C followed by staining with fixable viability stain (BD Horizon FVS700) for 10 minutes on ice in PBS. Subsequently, surface antigens were stained in FACS buffer for 30 minutes on ice. For cytokine staining, cells were then fixed and permeabilized by incubation in BD Fix/Perm solution for 15 minutes at room temperature followed by washing once in BD Perm/Wash solution. Intracellular cytokines were stained in BD Perm/Wash for 15 minutes at room temperature. For transcription factor staining, following surface staining, the cells were fixed using the Fixation/Permeabilization buffer in BD Transcription Factor kit for 45 minutes at 4°C. Transcription factors were stained in permeabilization buffer from the same kit for 45 minutes at 4°C. Conversely, for combined transcription factor and cytokine staining, after surface staining, the cells were fixed using the Fixation/Permeabilization buffer in FoxP3 transcription factor staining kit (eBiosciences) for 1 hour at 4°C. Cytokines and transcription factors were then stained in the permeabilization buffer from the same kit following the manufacturer’s guidelines.

All antibodies were used at a 1:200 dilution unless otherwise specified. Antibodies used herein were as follows: CD4-FITC (RM4-4,BD biosciences), CD19-FITC (6D5,Biolegend), TCRβ-APCeF780 (H57-597; eBioscience), TCRγδ-BV421 (GL3,BD biosciences), CD45-V500 (30-F11, BD biosciences), CD3-PECF594 (BM10-37, BD biosciences), RORγt-APC (B2D, BD biosciences), IL-17-BV786 (TC11-18H10, BD biosciences), IL-22-PE (1H8PWSR; eBioscience), T-bet-PECy7 (4B10,Biolegend), IFNγ-PerCP-Cy5.5 (XMG1.2; BD biosciences), CD69-V450 and Pe-CF594 (H1.2F3; BD biosciences), CCR6-Alexa Fluor 647 (140706; BD biosciences), CD27 PE-Cy7 (LG.3A10; BD biosciences), CD44-V500 (1M7; BD biosciences), Ki67-BV786 (B56; BD biosciences 1:100)

To determine the level ofpSTAT5, 1×10^6^ cells were fixed 100 µL with BD phosflow Lyse/Fix (diluted to 1x with water) for 10 minutes at 37°C. Subsequently, cells were washed once with FACS buffer and re-suspended in 100 µL BD phosflow perm buffer III, which was pre-chilled to −20°C, and incubated on ice for 30 minutes. Cells were then washed once with FACS buffer and stained for 30 minutes on ice in FACS buffer. Antibodies used in the staining of p-STAT5 were: CD4-FITC (GK1.5; BD Biosciences), CD8-APC 53-6.7; BD Biosciences) and pSTAT5-PEcy7 (47/Stat5 pY694; BD Biosciences; 5 µL/test). Samples were acquired using BD LSR Fortessa™ and BD FACSDiva software v8.0.2.

### Bone Marrow chimera

First, CD45.1^+^CD45.2^+^ mice were lethally irradiated (900 rad). Next day they were injected intravenously with 10×10^6^ cells of whole bone marrow cells from CD45.1^+^ wild-type and CD45.2^+^ RORγt^CRE^-STAT5^F/F^ mixed at 1:1 ratio. All Chimeras were analyzed after 12-14 weeks after reconstitution.

### Antibiotics administration

Pregnant female mice were treated with a cocktail of the following antibiotics: 1mg/ml Collistin, 5 mg/ml Streptomycin, 1mg/ml Ampicillin and 0.5mg/ml Vancomycin (all antibiotics Sigma) in their drinking water starting three days before delivery and until weaning of pups (3 weeks after the birth). The antibiotic-containing drinking water was replaced once a week until analysis.

### Citrobacter infection

*Citrobacter. rodentium* strain DBS100 (ATCC 51459; American Type Culture Collection) was purchased from ATCC and was cultured in Luria–Bertani broth overnight. CFU/ml (Dose) was determined by measuring the OD at 600 nm. Pups that were 10-12 days old were infected by oral gavage of 5×10^6^ CFU/mouse in a volume of 50µl. At day 6 post infection, the pups were euthanized and colons and fecal samples were collected. Bacterial load in the feces was determined as described (Sagaidak et al., 2016).

### Cell sorting, RNA extraction, cDNA synthesis and Real-Time PCR

Stained cells from thymi of 1-day old pups were sorted using BD ARIA FUSION™ and BD FACSDiva software v8.0.2. Target populations were sorted directly in RLT buffer (Qiagen) supplemented with 2-mercaptoethanol. Total RNA was extracted using Rneasy micro kit (Qiagen) and then used for cDNA synthesis using iScript cDNA synthesis kit (Biorad), according to the manufacturer’s protocol. SsoFast EvaGreen supermix (Biorad) was used to catalyze real-time PCR reactions, which were run on CFX96 (Biorad) and analyzed using Bio-rad CFX manager software. Gene expression levels were normalized to that of beta-actin. The following primers were used: *Actb*, Fwd-GGCTGTATTCCCCTCCATCG, Rev-CCAGTTGGTAACAATGCCATGT; *Stat5a*, Fwd-TCCGCAGCACCAGGTAAA, Rev-GGGATTATCCAAGTCAATAGCATC; *Stat5b*, Fwd-ACAACGGCAGCTCTCCAG, Rev-TGGGCAAACTGAGCTTGGATC.

### Data Analysis

Flow cytometry data was analyzed using Flow Jo V 10 software. All the statistical analyses and graphs were generated using Prism v7. All statistical tests used are described in the Figure legends

