## supplementary figures for "The neonatal microenvironment programs conventional and intestinal Tbet^+^ γδT17 cells through the transcription factor STAT5"

### Supplementary Figures and Legends

#### Figure S1

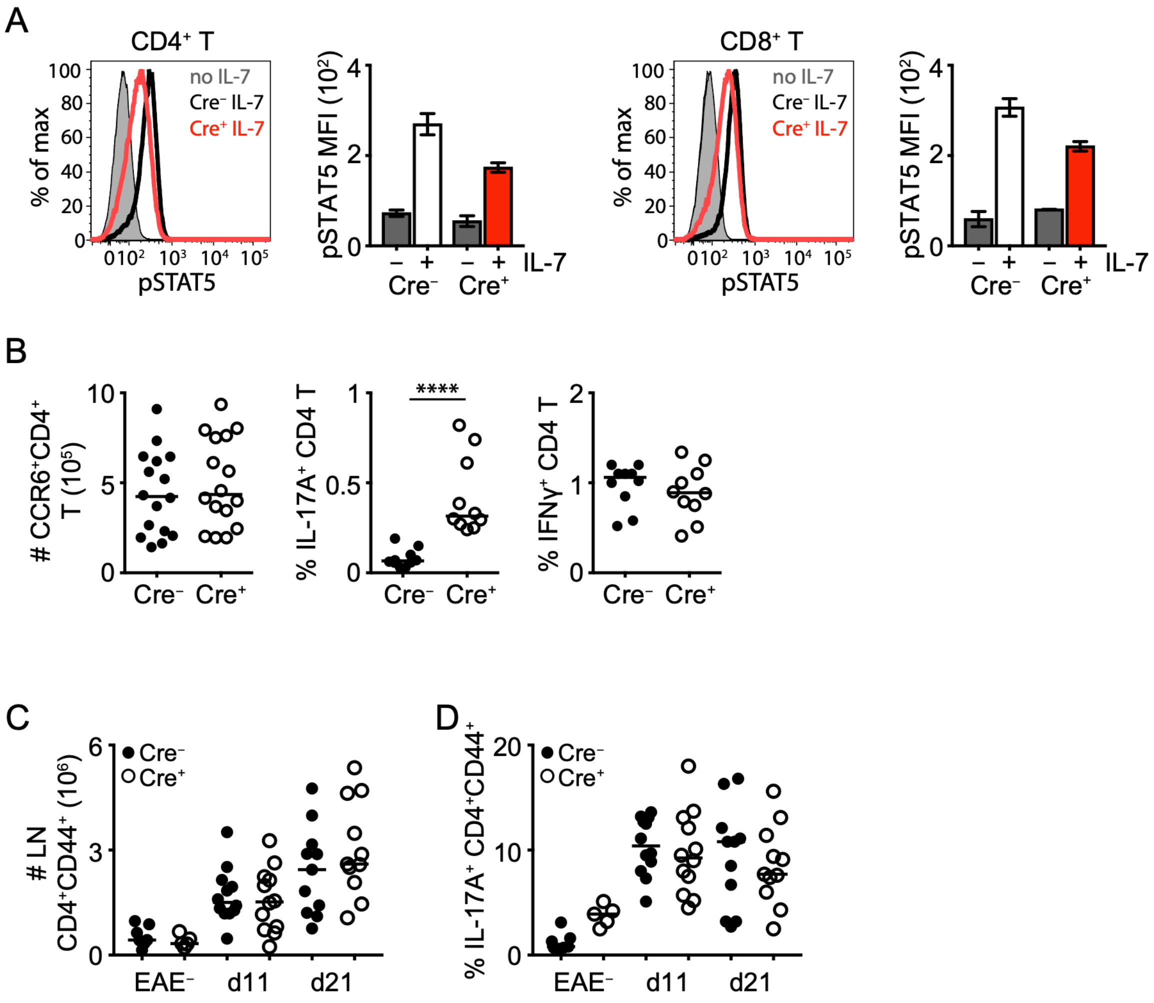


**Figure S1. CD4$^{+}$ and CD8$^{+}$ T cells in ROR$\gamma$t$^{CRE}$-STAT5$^{F/F}$ mice**

Flow cytometric analysis of LN CD4$^{+}$ and CD8$^{+}$ T cells in ROR$\gamma$t$^{CRE}$-STAT5$^{F/F}$ (Cre$^{+}$) and littermate control mice (Cre$^{-}$). In graphs each symbol represents a mouse and line the median. Bar graphs show median with range. *****p < 0.0001 using Mann-Whitney test.

(A) Expression of pSTAT5 (histograms) and pSTAT5 mean fluorescent intensity (MFI) (bar graphs) in CD4$^{+}$ (left) and CD8$^{+}$ T cells (right) following a 30 minute stimulation with 20 ng/ml recombinant murine IL-7 (representative of 2 experiments).

(B) Numbers of CD4$^{+}$CCR6$^{+}$ T cells (left), frequency of IL-17A$^{+}$ CD4$^{+}$ T cells (middle) and frequency of IFNγ$^{+}$ CD4$^{+}$ T cells (right).

(C-D) Numbers of $CD4^{+}CD44^{+}$ T cells (C) and frequency of IL-$17A^{+}$ $CD4^{+}CD44^{+}$ T cells (D) in the LNs of unimmunized control mice ($EAE^{-}$) and mice at days 11 and 21 after MOG-CFA immunization.

**Figure S2**

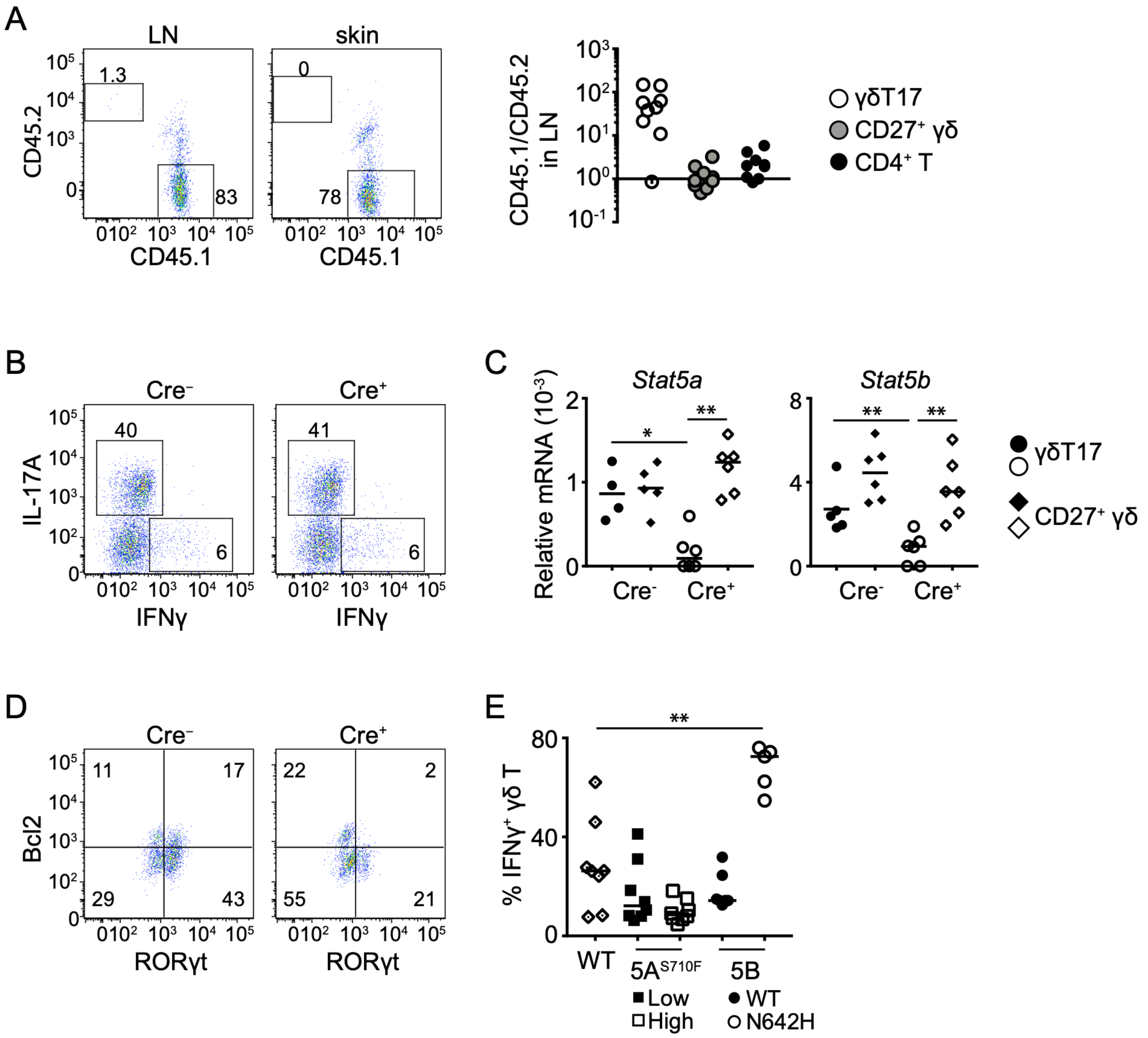


**Figure S2. Impact of STAT5 in γδT17 cells**

Flow cytometric analysis of γδ T cells in RORγt$^{CRE}$-STAT5$^{F/F}$ (Cre$^{+}$) and littermate control mice (Cre$^{-}$) or in STAT5A and STAT5B hyperactive mutant mice as described in Figure 2. In graphs, each symbol represents a mouse and line the median. *p < 0.05, **p < 0.01, ***p < 0.001, using Mann-Whitney test.

(A) Dot plots: frequency of CD45.1$^{+}$ (wild-type) and CD45.2$^{+}$ (STAT5-deficient) γδT17 cells from mixed BM chimera hosts (CD45.1$^{+}$CD45.2$^{+}$) in the LN and skin; graph shows the ratio of CD45.1$^{+}$/CD45.2$^{+}$ cells in the LN (each symbol represents a mouse).

(B) Expression of IL-17A and IFNγ in thymic γδ T cells one day after birth (representative of two litters with 2-5 pups per genotype per litter).

(C) Expression of *Stat5a* and *Stat5b* in sorted thymic CD27$^{-}$CD44$^{+}$CCR6$^{+}$ (γδT17) and CD27$^{+}$ γδ T cells one day after birth.

(D) Expression of BCL2 and RORγt in γδ T cells from LN of day 7 old mice (numbers indicate percent of expression; representative of two litters with 3-5 pups per genotype per litter).

(E) Expression of IFNγ within the LN γδ T cell compartment of the indicated STAT5A and STAT5B hyperactive mutant mice or WT control mice.

**Figure S3**

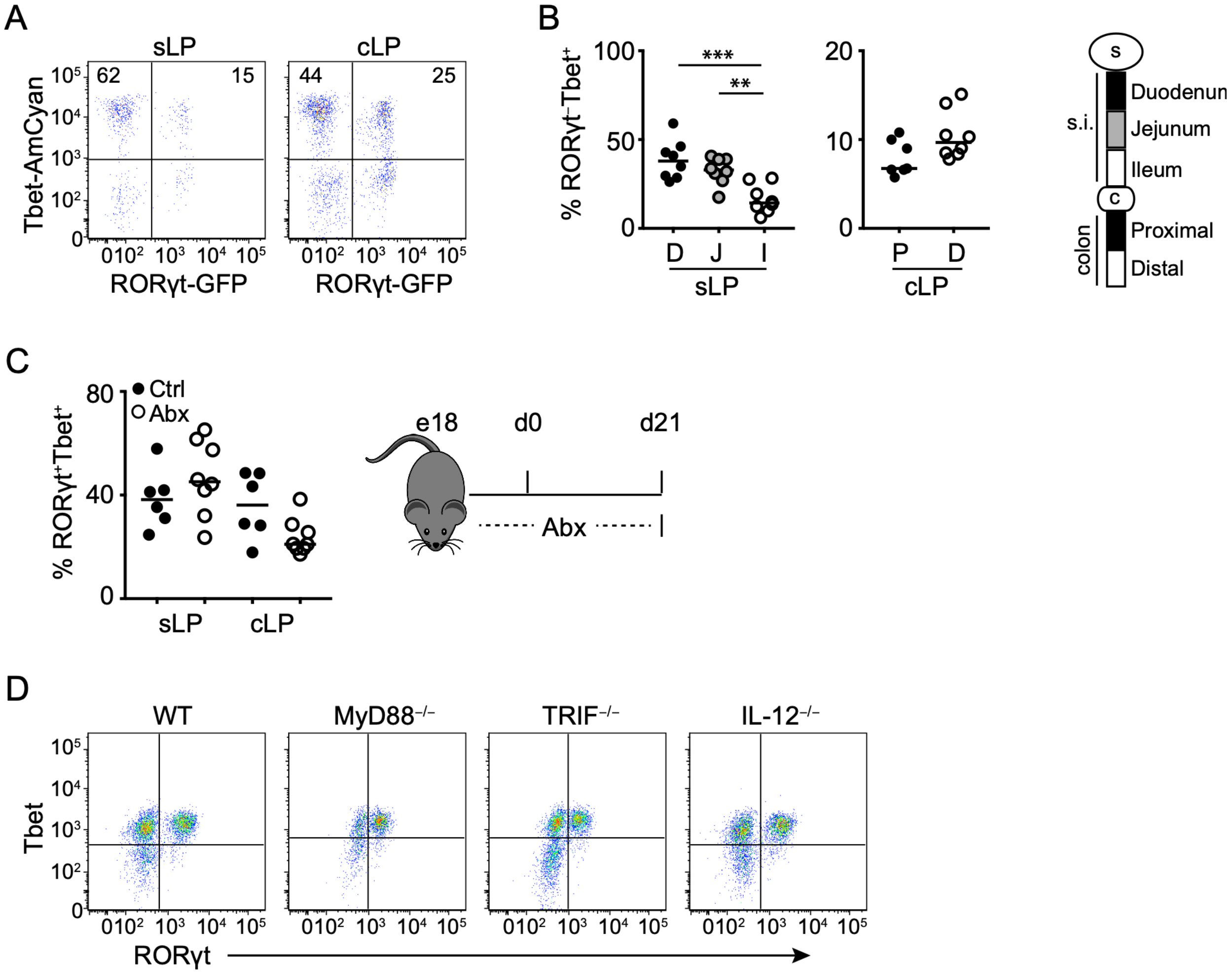


**Figure S3. Characterization of a novel Tbet$^{+}$ γδT17 population in the gut**

Flow cytometric analysis of intestinal γδ T cells from the indicated treatments and mouse strains. In graphs each symbol represents a mouse and line the median. *p < 0.05, **p < 0.01, ***p < 0.001, using Mann-Whitney test.

(A) Expression of AmyCyan and GFP in cLP and sLP γδ T cells from double transgenic mice reporting AmCyan under the control of the Tbet promoter and GFP under the control of the RORγt promoter (representative plots of 4 experiments).

(B) Frequency of RORγt$^{-}$Tbet$^{+}$ cells within the γδ T cell compartment in the indicated small intestinal and colonic segments.

(C) E18 pregnant mice were treated with an antibiotics (Abx) cocktail (see Methods) in the drinking water until their pups were analyzed at 21 days old. Graph shows the frequency of $ROR\gamma t^{+}Tbet^{+}$ cells within the sLP and cLP γδ T cell compartment in these pups.

(D) Expression of RORγt and Tbet in the cLP γδ T cell compartment in the indicated knockout mouse strains.

**Figure S4**

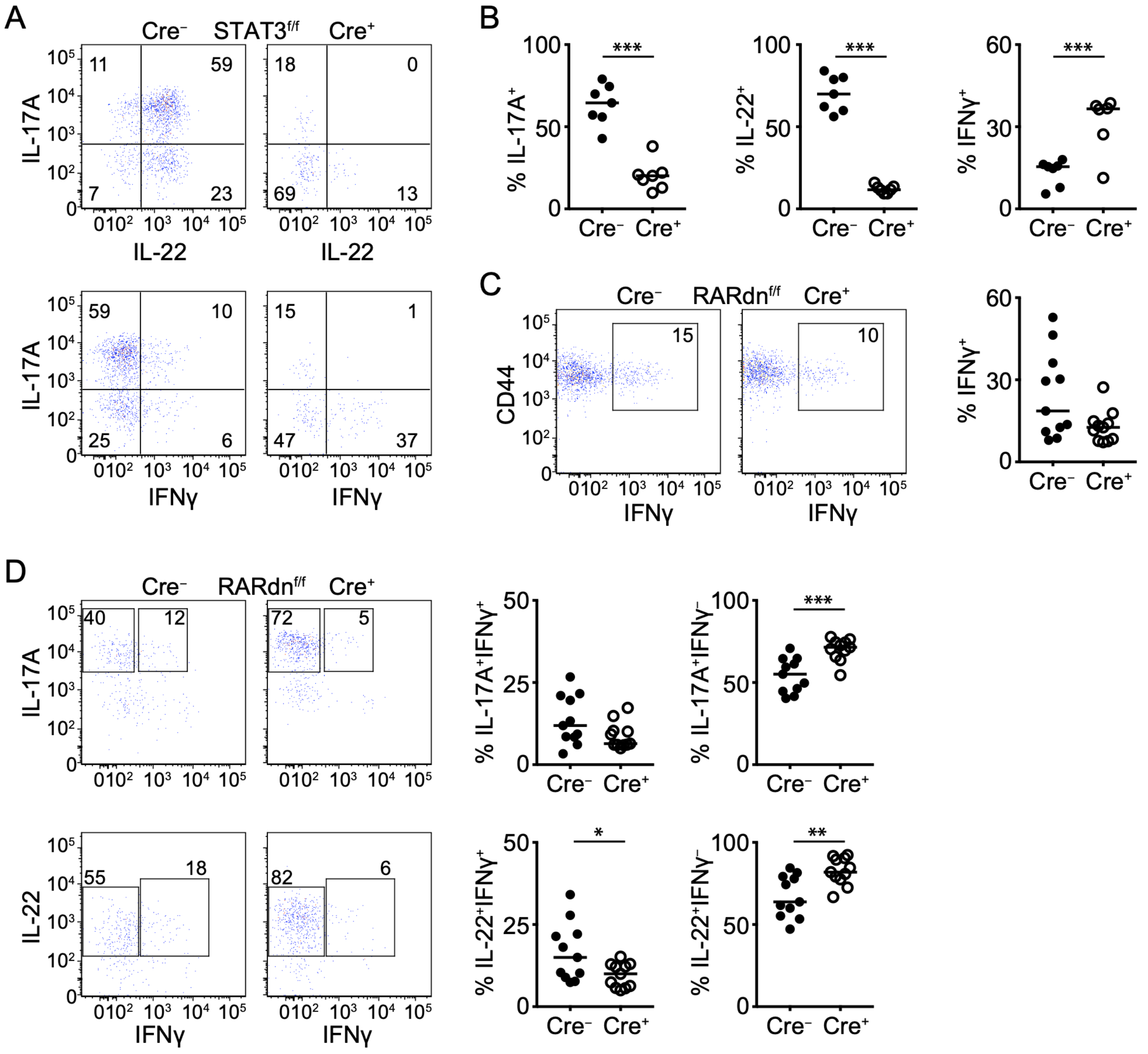


**Figure S4. STAT3 and retinoic acid receptor signaling regulate cytokine production in intestinal γδT17 cells**

Flow cytometric analysis of small intestinal γδ T cells in RORγt$^{CRE}$-STAT3$^{F/F}$ or RORγt$^{CRE}$-RARdn$^{F/F}$ (Cre$^{+}$) and littermate control mice (Cre$^{-}$) following IL-23 stimulation. In graphs each symbol represents a mouse and line the median. *p < 0.05, **p < 0.01, ***p < 0.001 using Mann-Whitney test.

(A) Expression of IL-17A and IL-22 (top) or IL-17A and IFNγ (bottom) in RORγt$^{CRE}$-STAT3$^{F/F}$ (Cre$^{+}$) and littermate control mice (Cre$^{-}$). Numbers indicate percent of positive expression.

(B) Frequency of IL-17A$^{+}$, IL-22$^{+}$ and IFNγ$^{+}$ γδ T cells in RORγt$^{CRE}$-STAT3$^{F/F}$ (Cre$^{+}$) and littermate control mice (Cre$^{-}$).

(C) Expression of CD44 and IFNγ (dot plots) and frequency of IFNγ$^{+}$ γδ T cells (graph) in RORγt$^{CRE}$-RARdn$^{F/F}$ (Cre$^{+}$) and littermate control mice (Cre$^{-}$).

(D) Expression of IL-17A and IFNγ (top dot plots) or IL-22 and IFNγ (bottom dot plots) with graphical representation of the frequency of IL-17A$^{+}$IFNγ$^{+}$ and IL-17A$^{+}$IFNγ$^{-}$ or IL-22$^{+}$IFNγ$^{+}$ and IL-22$^{+}$IFNγ$^{-}$ γδ T cells in RORγt$^{CRE}$-RARdn$^{F/F}$ (Cre$^{+}$) and littermate control mice (Cre$^{-}$).

**Figure S5**

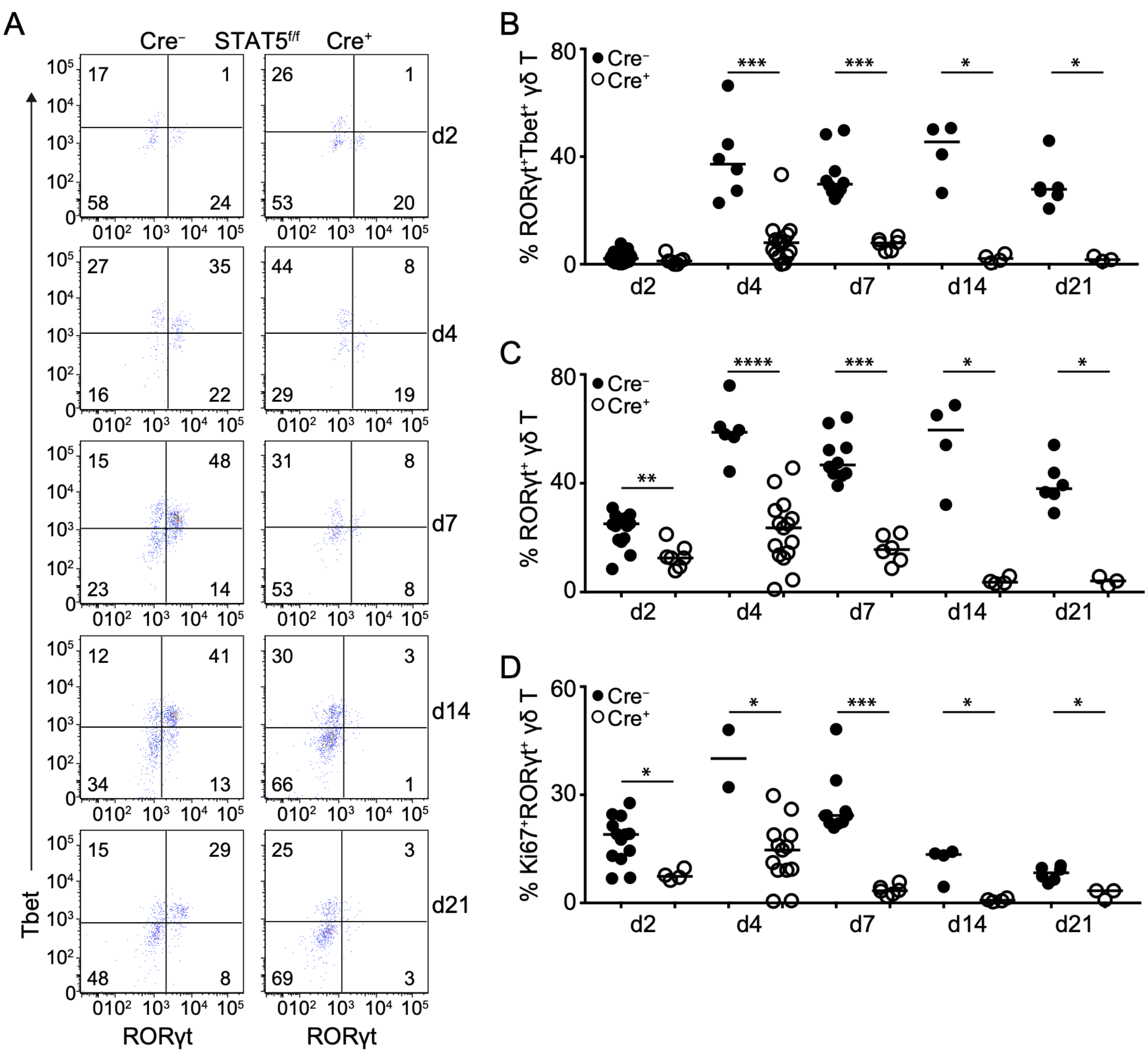


**Figure S5. STAT5 regulates the neonatal fate of intestinal Tbet⁺ γδT17 cells**

Flow cytometric analysis of small intestinal γδ T cells in RORγt$^{CRE}$-STAT5$^{F/F}$ (Cre$^{+}$) and littermate control mice (Cre$^{-}$) during neonatal ontogeny. Day of birth is counted as day(d)1. In graphs each symbol represents a mouse and line the median. *$p < 0.05$, **$p < 0.01$, ***$p < 0.001$, ****$p < 0.0001$ using Mann-Whitney test.

(A) Expression of RORγt and Tbet within the γδ T cell compartment of sLP at the indicated days after birth. Numbers indicate percent of RORγt and Tbet expression.

(B) Frequency of sLP RORγt$^{+}$Tbet$^{+}$ γδ T cells at the indicated days after birth.

(C) Frequency of sLP RORγt$^{+}$ γδ T cells (including Tbet$^{+}$) at the indicated days after birth.

(D) Frequency of sLP Ki67$^{+}$RORγt$^{+}$ γδ T cells at the indicated days after birth.

**Figure S6**

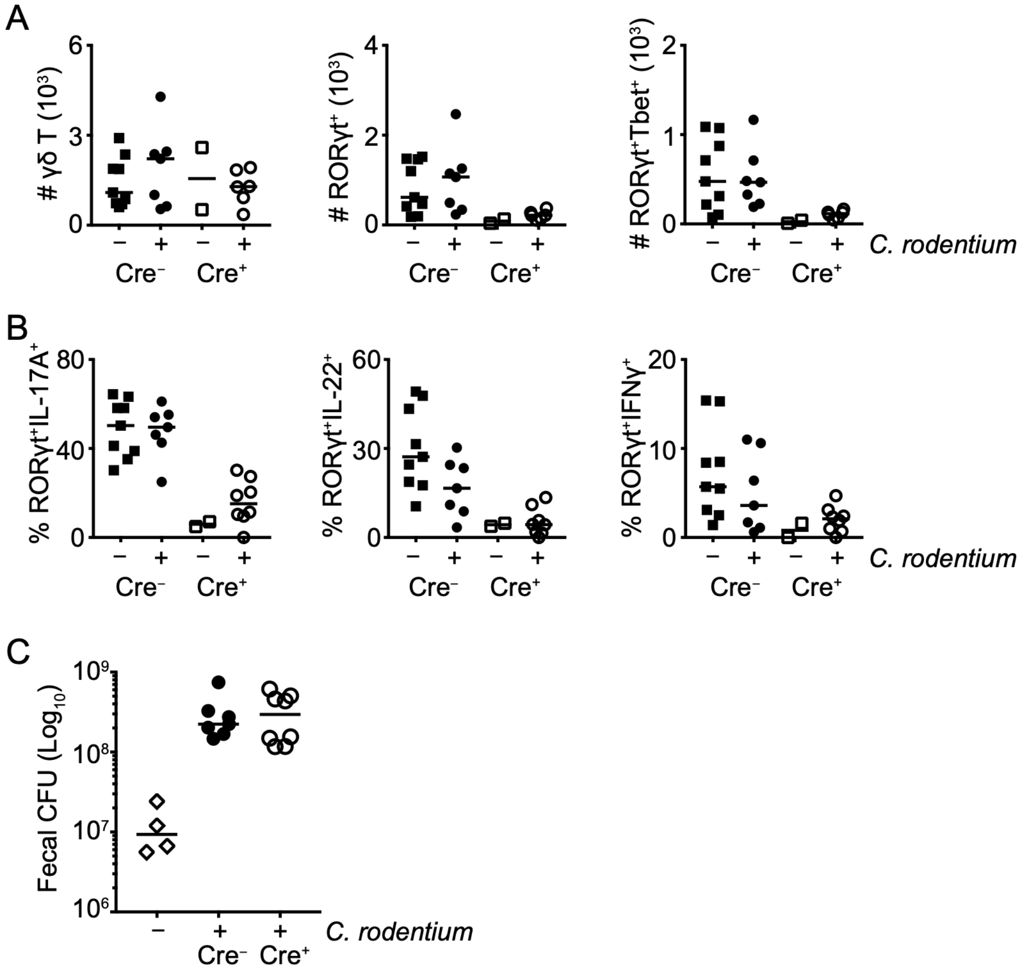


**Figure S6. Intestinal γδT17 cells are not required for neonatal bacterial infection**

Flow cytometric analysis of cLP γδ T cells in RORγt$^{CRE}$-STAT5$^{F/F}$ (Cre$^{+}$) and littermate control mice (Cre$^{-}$) before or after infection with *Citrobacter rodentium*. In graphs, each symbol represents a mouse and line the median. Cytokine detection was performed following IL-23 re-stimulation.

(A-B) Day 10-12 old pups were infected orally with C. *rodentium* and cLP cells were analyzed 6 days later. (A) Numbers of total (left), RORγt$^{+}$ (middle) and RORγt$^{+}$Tbet$^{+}$ (right) γδ T cells in infected and uninfected mice. (B) Frequency of RORγt$^{+}$IL-17A$^{+}$ (left), RORγt$^{+}$IL-22$^{+}$ (middle) and RORγt$^{+}$IFNγ $^{+}$ (right) γδ T cells in infected and uninfected mice.

(C) Fecal colony forming units (CFU) in infected and uninfected mice detected by qPCR.
